# Sensitization of Human and Rat Nociceptors by Low Dose Morphine is TLR4-dependent

**DOI:** 10.1101/2023.12.19.572472

**Authors:** Eugen V. Khomula, Jon D. Levine

**Affiliations:** Department of Oral & Maxillofacial Surgery, and Division of Neuroscience, University of California at San Francisco, 513 Parnassus Avenue, San Francisco, CA 94143, USA; Departments of Medicine and Oral & Maxillofacial Surgery, and Division of Neuroscience, UCSF Pain and Addiction Research Center, University of California at San Francisco, 513 Parnassus Avenue, San Francisco, CA 94143, USA

**Keywords:** morphine, Toll-like Receptor 4 (TLR4), opioid-induced hyperalgesia (OIH), human and rat nociceptors, sensitization, excitability, patch-clamp electrophysiology

## Abstract

While opioids remain amongst the most effective treatments for moderate-to-severe pain, their substantial side effect profile remains a major limitation to broader clinical use. One such side effect is opioid-induced hyperalgesia (OIH), which includes a transition from opioid-induced analgesia to pain enhancement. Evidence in rodents supports the suggestion that OIH may be produced by the action of opioids at Toll-like Receptor 4 (TLR4) either on immune cells that, in turn, produce pronociceptive mediators to act on nociceptors, or by a direct action at nociceptor TLR4. And, sub-analgesic doses of several opioids have been shown to induce hyperalgesia in rodents by their action as TLR4 agonists. In the present *in vitro* patch-clamp electrophysiology experiments, we demonstrate that low dose morphine directly sensitizes human as well as rodent dorsal root ganglion (DRG) neurons, an effect of this opioid analgesic that is antagonized by LPS-RS Ultrapure, a selective TLR4 antagonist. We found that morphine (100 nM) reduced rheobase in human (by 36%) and rat (by 26%) putative C-type nociceptors, an effect of morphine that was markedly attenuated by preincubation with LPS-RS Ultrapure. Our findings support the suggestion that in humans, as well as in rodents, OIH is mediated by the direct action of opioids at TLR4 on nociceptors.

## Introduction

Chronic pain affects roughly 20% of adults ^20^, impacting more than 1 billion people globally. While opioid analgesics remain amongst the most potent and effective treatments for moderate-to-severe pain their side-effects, including loss of analgesic efficacy and worsening of pain (opioid-induced hyperalgesia, OIH), remain major clinical limitations, especially for their use to treat chronic pain. It is well-established that opioids produce analgesia by action at their cognate receptors, predominantly the mu-opioid receptor (MOR) ^45^. However, some side effects are thought to be mediated by the action of opioid analgesics at other receptors, most well characterized for Toll-like receptor 4 (TLR4). Thus, several opioid analgesics (e.g. morphine, fentanyl, remifentanil, and oxycodone) bind to and activate TLR4 even at low, sub-analgesic concentrations ^13, 24, 25^. And, the systemic administration of sub-analgesic doses of opioid analgesics can, paradoxically, produce hyperalgesia (OIH) ^22^, an effect that is mediated either directly, by activation of TLR4, which is present on nociceptors ^40, 53^, or indirectly by the action of opioids at TLR4 on non-neuronal cells, including cells of the immune system that, in turn, release pronociceptive mediators ^37, 47^. The primary afferent nociceptor has been suggested to play a key role in OIH ^9^, which is attenuated by TLR4 antagonists ^38^, as well as by intrathecal administration of an oligodeoxynucleotide (ODN) antisense for TLR4 mRNA ^2^. Therefore, while opioids act as MOR agonists, to produce analgesia that is, in part, dependent on nociceptor MORs ^2, 16^, tolerance to MOR-dependent analgesia could unmask a hyperalgesic action of opioids, mediated by TLR4. To date, our understanding of the role of TLR4 in the side-effects of opioid analgesics derives from data obtained from rodents. And, while TLR4 is present in human as well as in rodent dorsal root ganglion (DRG) neurons ^5, 35, 36, 53^, differential expression of genes affecting DRG function may be responsible for species specific divergence in nociceptor functions ^28^. We have previously shown, in the rat, that higher doses of morphine (starting at ∼1 mg/kg) induces analgesia that is dependent on nociceptor MOR, while low, sub-analgesic doses of morphine (3-30 μg/kg) induce hyperalgesia by action at nociceptors TLR4 ^2, 16^. However, it still remains to be established if opioids can similarly, act at TLR4 in human DRG neurons, to contribute to OIH. To address this question, in the present experiments we used *in vitro* patch-clamp electrophysiology to evaluate for an increase in excitability of putative nociceptors, induced by acute administration of morphine (100 nM), a concentration estimated to be produced by a single sub-analgesic dose, 0.15 mg/kg ^44^, using cultured human and rat DRG neurons, to establish parallel *in vitro* models of OIH.

We report that the low concentration of morphine (100 nM) similarly sensitizes human as well as rat nociceptors, *in vitro*, an effect that is prevented by the highly selective TLR4 antagonist, LPS-RS Ultrapure ^33^, in both species. Our findings support the suggestion that in humans, as in rodents, OIH is mediated by the action of opioid analgesics at TLR4 on primary afferent nociceptors.

## Materials and Methods

### Culturing rat DRG neurons

Primary cultures of dorsal root ganglia (DRG) were made from 220 – 235 g adult male Sprague-Dawley rats, as described previously ^3, 17, 18, 31^. Under isoflurane anesthesia, rats were decapitated, and the dorsum of their vertebral column surgically removed; L_4_ and L_5_ DRGs were rapidly extracted, bilaterally, chilled and desheathed in Hanks’ balanced salt solution (HBSS), on ice. DRG were then treated with 0.25% collagenase Type 4 (Worthington Biochemical Corporation, Lakewood, NJ, USA) in HBSS for 18 minutes at 37°C, and then with 0.25% trypsin (Worthington Biochemical Corporation) in calcium- and magnesium-free PBS (Invitrogen Life Technologies, Grand Island, NY USA) for 6 minutes, followed by three washes and trituration in Neurobasal-A medium (Invitrogen Life Technologies) to produce a single-cell suspension. This suspension was centrifuged at 1000 RPM for 3 minutes followed by re-suspension in Neurobasal-A medium that was supplemented with 50 ng/mL nerve growth factor, 100 U/mL penicillin/streptomycin, B-27, GlutaMAX and 10% FBS (Invitrogen Life Technologies). Cells were then plated on cover slips and incubated at 37°C in 3.5% CO_2_ for at least 24 hours before they were used in electrophysiology experiments.

### Culturing human DRG neurons

Human DRGs from a single male donor were purchased from Anabios (San Diego, CA, USA). Dissociated human DRGs from this donor were shipped to UCSF as a suspension in cryoprotective gel, on ice. They were delivered within hours after preparation of the suspension from donor tissue at the Anabios facility. All samples underwent visual inspection for tissue integrity in our laboratory and in parallel at Anabios, using fluorescent microscopy and electrophysiology. Human tissue originating from AnaBios was obtained with legal consent from US-based organ donors, adhering to United Network for Organ Sharing (UNOS) donor screening and consent standards. AnaBios follows procedures endorsed by the US Centers for Disease Control (CDC), subject to biannual inspections by the US Department of Health and Human Services (DHHS). Distribution of tissue to investigators, by Anabios, is governed by internal IRB procedures and is in compliance with Health Insurance Portability and Accountability Act (HIPAA) regulations. All organ transfers to AnaBios are traceable and periodically reviewed by US Federal authorities, ensuring the privacy of donor information during studies involving human DRGs. Thus, no identifying information was shared with our research team except that the organ donor from whom we received DRG had not succumbed to an opioid overdose.

Pursuant to an Anabios protocol, upon arrival at UCSF (San Francisco, CA, USA) the content of the transportation vial (2 mL of protective media, containing ∼1000 neurons) was transferred to 9 mL of wash media (DMEM/F12 with penicillin/streptomycin [Lonza; Allendale, NJ]), then recovered by centrifugation (∼350 xG ) at room temperature for 3 min, and resuspended in 0.6 mL of culturing media provided by Anabios, made on the basis of DMEM/F12 with penicillin/streptomycin and supplements: 10% horse serum (Thermo Fisher Scientific, Rockford, IL, USA), 2 mM glutamine, 25 ng/mL hNGF (Cell Signaling Technology, Danvers, MA, USA) and 25 ng/mL GDNF (PeproTech, Rocky Hill, NJ, USA) ^11^.

Cells from dissociated DRG were plated on 3 round (15 mm diameter) glass coverslips (0.2 mL of suspension per coverslip, in separate Petri dishes) that had been precoated with poly-D-lysine (reported by Anabios to be important for survival) (Neuvitro Corp., Camas, WA, USA). One hour after plating, 1 mL of the culture media was gently added and coverslips fractured such that 6-8 individual pieces of a coverslip that could be used for electrophysiology recordings was obtained. As recommended by Anabios, human DRG neurons were used after incubating for 48 h (5% CO2, 37°C), for up to day 10. Culture media in Petri dishes was exchanged every 2 days by replacing half of the media.

### Whole-cell patch-clamp electrophysiology

Following placement of individual coverslip fragments, plated with cells from dissociated human or rat DRGs, in the recording chamber, culture medium was substituted with the solution used to perform electrophysiology, Tyrode’s solution containing 140 mm NaCl, 4 mm KCl, 2 mm MgCl_2_, 2 mm CaCl_2_, 10 mm glucose, and 10 mm HEPES, adjusted to pH 7.4 with NaOH, with an osmolarity of 310 mOsm/kg. The recording chamber had a volume of 150 µl, and its perfusion system had a flow rate of 0.5–1 ml/min. Electrophysiology experiments were conducted at room temperature (20– 23°C).

Cells were identified as neurons by their double birefringent plasma membranes ^8,34^. Whole-cell patch-clamp recordings, performed in current clamp mode, were used to evaluate for changes in the excitability of cultured rat and human DRG neurons. Holding current was adjusted to maintain membrane potential at −70 mV. Rheobase, defined as the minimum magnitude of a current step needed to elicit an action potential (AP), was determined through a testing protocol utilizing a series of square wave pulses with current magnitude increasing by a constant step every sweep, until an AP was elicited. An initial estimate of rheobase was made with 500-pA increments (0.5 – 4 nA). The increments were then adjusted to achieve 5–10% precision of the rheobase estimate ^18, 30^.

Recording electrodes were fashioned from borosilicate glass capillaries (0.84/1.5 mm i.d./o.d., Warner Instruments, LLC) using a Flaming/Brown P-87 microelectrode puller (Sutter Instrument Co, Novato, CA, USA). After being filled with a solution containing 130 mm KCl, 10 mm HEPES, 10 mm EGTA, 1 mm CaCl_2_, 5 mm MgATP, and 1 mm Na-GTP; pH 7.2 (adjusted with Tris-base), resulting in 300 mOsmol/kg osmolarity, the recording electrode resistance was approximately 2 MΩ. Junction potential was not adjusted, and series resistance was below 10 MΩ at the end of recordings, without compensation. Recordings were conducted using an Axon MultiClamp 700 B amplifier, filtered at 20 kHz, and sampled at 50 kHz through an Axon Digidata 1550B controlled by pCLAMP 11 software (all from Molecular Devices LLC, San Jose, CA, USA).

To ensure the stability of baseline current, drugs were applied at least 5 minutes after the establishment of whole-cell configuration.

### Drugs and media

The following drugs were used in this study: Morphine sulfate salt pentahydrate, NaCl, KCl, MgCl_2_, CaCl_2_, NaOH, MgATP, Na-GTP, D-Glucose, 4-(2-Hydroxyethyl)piperazine-1-ethanesulfonic acid (HEPES), Ethylene glycol-bis(2-aminoethylether)-N,N,N′,N′-tetraacetic acid (EGTA) (Sigma Aldrich, St. Louis, MO, USA), LPS-RS Ultrapure (a selective TLR4 antagonist, lipopolysaccharide from *Rhodobacter* 9 *sphaeroides*, purchased from InvivoGen, San Diego, CA, USA), Collagenase Type 4, Trypsin (Worthington Biochemical Corporation, Lakewood, NJ, USA), Calcium- and Magnesium-free HBSS, Calcium- and Magnesium-free PBS, Neurobasal-A medium, B-27 supplement, GlutaMAX, Fetal Bovine Serum (FBS) (Invitrogen Life Technologies, Grand Island, NY USA), rat (recombinant) nerve growth factor (NGF)-beta, glutamine (Sigma Aldrich), penicillin/streptomycin, DMEM/F12 with penicillin/streptomycin (Lonza, Allendale, NJ, USA), horse serum (Thermo Fisher Scientific, Rockford, IL, USA), human NGF (hNGF, Cell Signaling Technology, Danvers, MA, USA), human GDNF (Peprotech, Rocky Hill, NJ, USA).

Stock solution of LPS-RS Ultrapure was prepared in purified water (1 mg/mL) and stored at -20°C in 50 µL vials. One vial was thawed on the day of the experiment and stored at 4°C. Final concentration of LPS-RS Ultrapure (10 µg/mL) was achieved by 1:99 dilution in Tyrode’s solution, performed just before it was used in experiments.

Stock solution of morphine sulfate (0.5 mM) was prepared from powder (0.76 mg/mL, in purified water) freshly on the day of the experiment and stored at 4°C. The final concentration of morphine in perfusion solution was selected to be 100 nM (76 ng/mL; 1:4999 dilution of the stock solution, performed in two steps, 1:99 in purified water then 1:49 in Tyrode’s solution), which we consider as a low concentration, as it was 10-100x lower than typical concentration used in most *in vitro* studies (1-10 µM) ^6, 12, 50, 56^. We estimated that plasma concentration of 100 nM would be produced by a single intravenous sub-analgesic dose (0.15 mg/kg, i.e. 10 mg/70kg) based on the reported findings that, on average, a dose of 3.0 mg of morphine resulted in plasma level of 40.2 ng/ml in human subjects who had an average weight of 56.3 kg (Table 1 and 2 from ^44^). We have previously shown that, in the rat, when performing a dose response curve, morphine induces analgesia at doses starting at ∼1 mg/kg ^16^.

### Statistical analysis

Rheobase was measured before and 5 min after application of morphine. Magnitude of the effect of morphine was expressed as percentage reduction in rheobase (i.e., value before morphine administration was subtracted from value after, then the difference was divided by pre drug baseline). The following statistical tests were used: one-sample two-tailed Student’s *t*-test versus zero and two-sample unpaired two-tailed Student’s *t*-test.

Prism 10.1 (GraphPad Software) was used to generate graphics and to perform statistical analyses; *P<0.05* is considered statistically significant. Data are presented as mean ± SEM.

## Results

### Nociceptor sensitization by low dose morphine

To test the hypothesis that low dose morphine sensitizes small-diameter human and rat DRG neurons, we performed *in vitro* patch-clamp electrophysiology, evaluating the effect of our low concentration of morphine (100 nM, i.e. 76 ng/mL, corresponding to a single sub-analgesic dose of 0.15 mg/kg) ^16, 44^ on neuronal excitability. Rheobase, the minimal sustained current required to generate an action potential (AP), was selected for our measure of neuronal excitability as it is a well-defined electrophysiological property that reflects excitability in DRG neurons ^11, 29, 30, 51, 52^. Percentage decrease in rheobase was used as the measure of nociceptor sensitization. The effect of 100 nM morphine on rheobase was evaluated in putative C-type nociceptive (small-diameter) neurons from rat (soma diameter <30 µm) ^19, 21, 46, 54^ and human (soma diameter 40-60 µm) DRG ^7, 15, 58^. We found that low dose morphine induces a significant decrease in rheobase in small-diameter rat (26±7%, n=6, * p = 0.013, t_(5)_ = 3.8, Fig. 1B; illustrative traces in Fig. 1A) and human (36±7%, n=9, ** p = 0.0014, t_(8)_ = 4.8, Fig. 2B; illustrative traces in Fig. 2A) DRG neurons.

**Figure 1.**
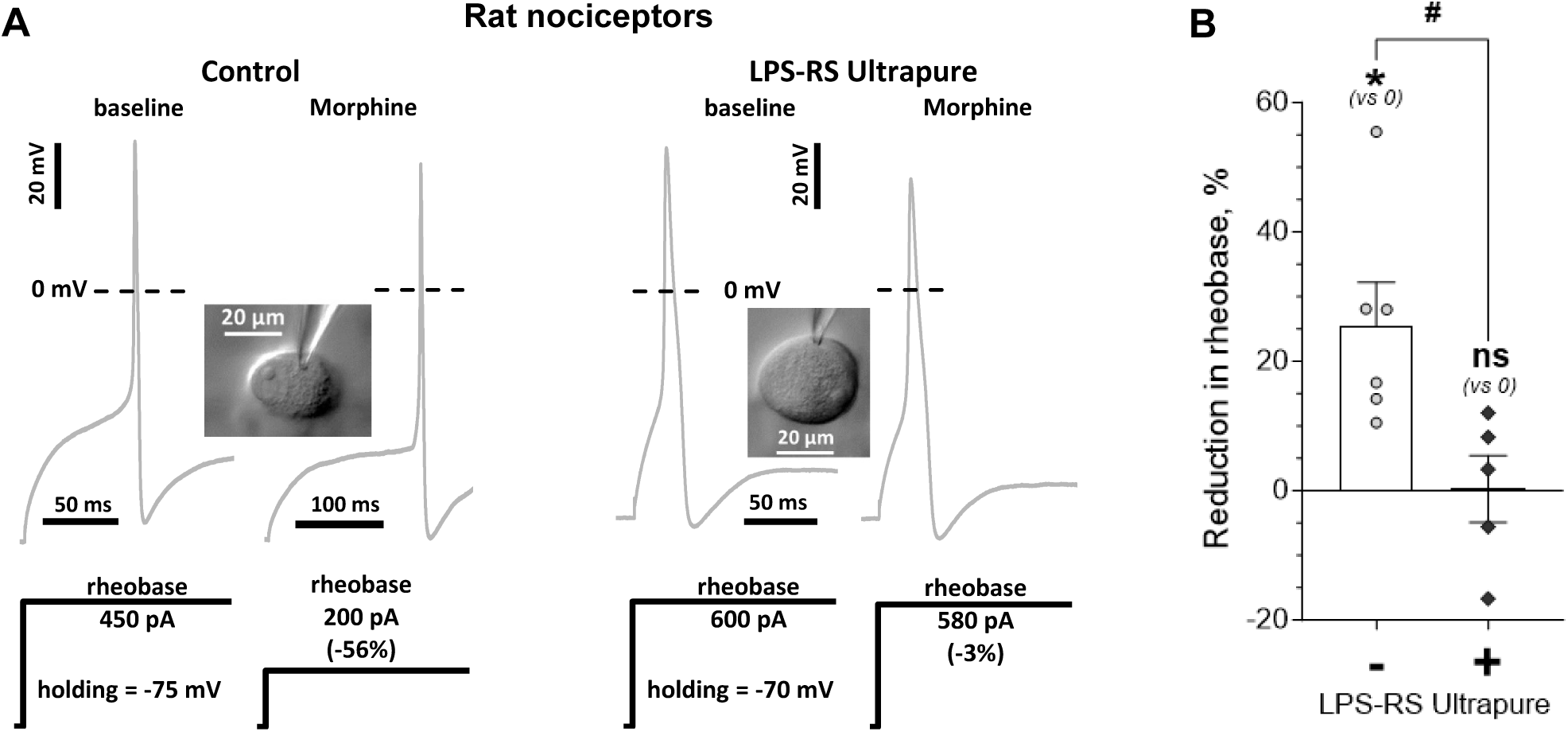
Low dose morphine induces TLR4-dependent sensitization of rat nociceptors. **A.** Examples of low dose morphine induced reduction in rheobase of putative C-type rat nociceptors, and its prevention by a selective TLR4 antagonist (LPS-RS Ultrapure). Electrophysiological traces (upper, grey) show APs generated in response to stimulation of a small-diameter DRG neuron (depicted in the inset image) with a square wave current pulse (shown below AP recordings, black). The height of the pulse represents rheobase. The scale is indicated by corresponding scale bars and, if not indicated by a different scale bar, is the same for left and right traces. Dotted line shows level of 0 mV. *Left panel* shows traces from a low dose morphine-treated neuron, of the control group, with no TLR4 antagonist added. Note the reduction in the height of the current pulse *(right traces)* after application of morphine (100 nM), compared to the baseline value *(left trace)*. *Right panel* shows traces from a neuron of the “prevention” protocol group, preincubated for 30 min with the selective TLR4 antagonist, LPS-RS Ultrapure (10 μg/mL). Note, only a small reduction in rheobase in response to morphine (100 nM), being markedly attenuated compared to the control effect of morphine in the absence of LPS-RS Ultrapure. **B.** Reduction in rheobase, relative to its baseline value, produced by morphine (100 nM) in rat small-diameter DRG neurons, without (control group; left white bar, “-“) and after preincubation with the selective TLR4 antagonist, LPS-RS Ultrapure (10 μg/mL; “prevention” protocol group; right grey bar, “+”). Bars show mean ± S.E.M. Symbols show effect in individual neurons. Morphine produced a significant reduction in rheobase in the control group (one-sample two-tailed Student’s *t*-test for zero effect: * p = 0.013, t_(5)_ = 3.8) that was significantly attenuated in the “prevention” group (two-sample unpaired two-tailed Student’s *t*-test: ^#^ p = 0.018, t_(9)_ = 2.9), became not significantly different from baseline (one-sample two-tailed Student’s *t*-test for zero effect: p = 0.96, t_(4)_ = 0.05), supporting the suggestion that sensitization produced by low dose of morphine, in rat nociceptors, is dependent on TLR4 that is expressed in the nociceptor. Number of cells: 6 in control, 5 in the “prevention” protocol group.

**Figure 2.**
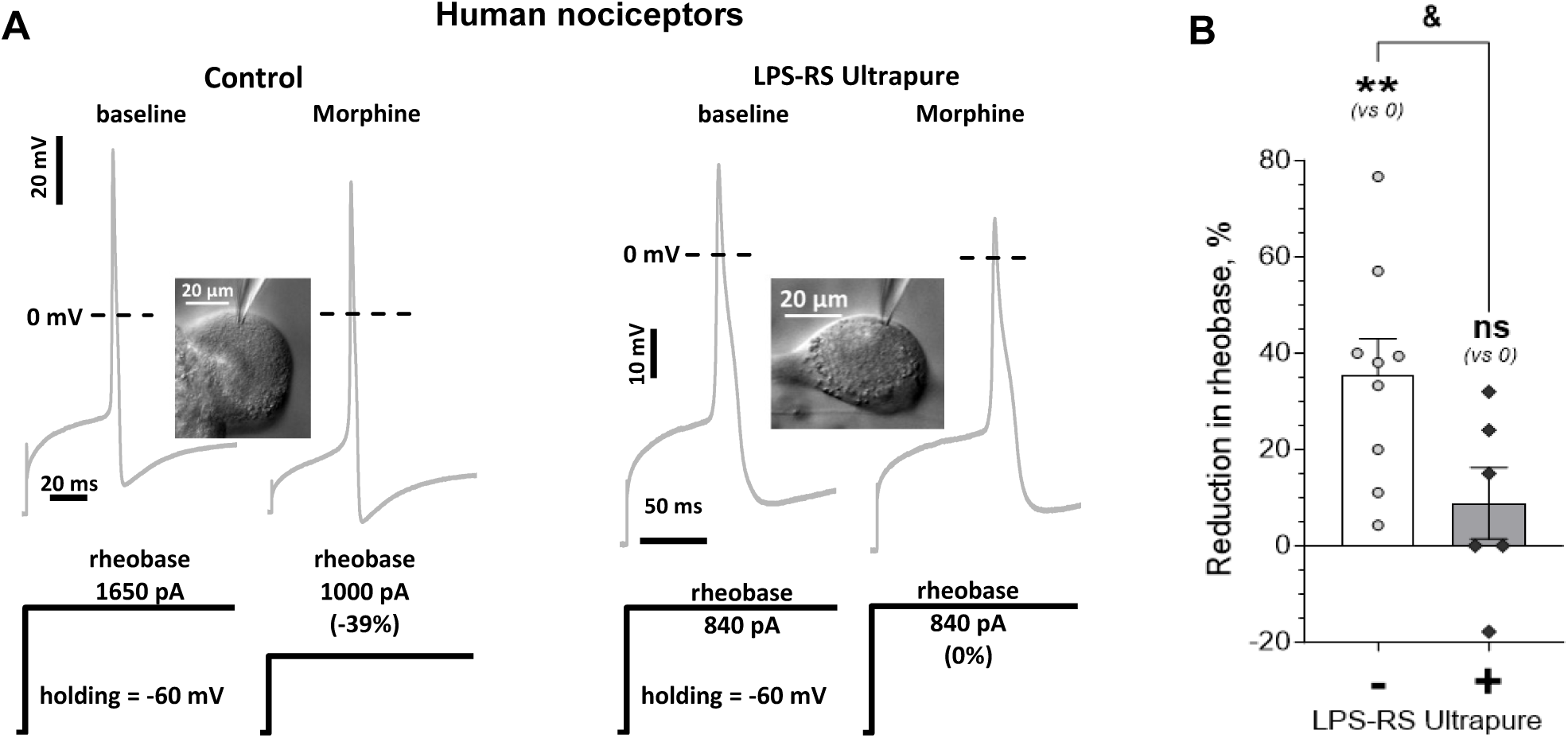
Low dose morphine induces TLR4-dependent sensitization of human nociceptors. **A.** Examples of low dose morphine induced reduction in rheobase in putative C-type human nociceptors, and its prevention by a selective TLR4 antagonist (LPS-RS Ultrapure). Electrophysiological traces (upper, grey) show APs generated in response to stimulation of a small-diameter DRG neuron (depicted in the inset image) with a square wave current pulse (shown below AP recordings, black). The height of the pulse represents rheobase. The scale is indicated by corresponding scale bars and, if not indicated by a different scale bar, is the same for left and right traces. Dotted line shows level of 0 mV. *Left panel* shows traces from a low dose morphine-treated neuron of the control group, with no TLR4 antagonist added. Note the reduction in the height of the current pulse *(right traces)* after application of morphine (100 nM) compared to baseline value *(left trace)*. *Right panel* shows traces from a neuron of the “prevention” protocol group, preincubated for 30 min with the selective TLR4 antagonist, LPS-RS Ultrapure (10 μg/mL). In this cell no reduction in rheobase in response to morphine (100 nM) is observed, being markedly attenuated compared to the control effect of morphine in the absence of LPS-RS Ultrapure. **B.** Reduction in rheobase, relative to its baseline value, produced by morphine (100 nM) in human small-diameter DRG neurons, without (control group; left white bar, “-“) and after preincubation with the selective TLR4 antagonist, LPS-RS Ultrapure (10 μg/mL; “prevention” protocol group; right grey bar, “+”). Bars show mean ± S.E.M. Symbols show effect in individual neurons. Morphine produced a significant reduction in rheobase in the control group (one-sample two-tailed Student’s *t*-test for zero effect: ** p = 0.0014, t_(8)_ = 4.8) that was significantly attenuated in the “prevention” group (two-sample unpaired two-tailed Student’s *t*-test: ^&^ p = 0.031, t_(13)_ = 2.4), became not significantly different from baseline (one-sample two-tailed Student’s *t*-test for zero effect: p = 0.29, t_(5)_ = 1.2), supporting the suggestion that sensitization produced by low dose of morphine, in human nociceptors, is dependent on TLR4 that is expressed in the nociceptor. Number of cells: 9 in control, 6 in the “prevention” protocol group.

### Prevention of the sensitization of nociceptors by low dose morphine with a selective TLR4 antagonist

We next evaluated the contribution of TLR4 to low dose morphine-induced sensitization of rat and human DRG neurons, *in vitro*. In both rat and human small-diameter DRG neurons that were pretreated with the selective TLR4 antagonist, LPS-RS Ultrapure (10 μg/mL, for 30 min before, during, and continued after the exposure to morphine), reduction in rheobase induced by low dose morphine (relative to baseline in the same cell) was significantly attenuated, compared to morphine treatment without the TLR4 antagonist (rat: ^#^ p = 0.018, t_(9)_ = 2.9; human: ^&^ p = 0.031, t_(13)_ = 2.4) and was not significantly different from zero (no effect) (rat: 0±5%, n=5, 95% CI = (-14 .. +15)%, p = 0.96, t_(4)_ = 0.05, Fig. 1B; illustrative traces in Fig. 1A; human: 9±7%, n=6, 95% CI = (-10.. +28)%, p = 0.29, t_(5)_ = 1.2, Fig. 2B; illustrative traces in Fig. 2A).

## Discussion

Opioid-induced hyperalgesia (OIH) remains a paradoxical effect of several clinically used opioid analgesics (e.g., morphine, fentanyl, remifentanil and oxycodone). It has long been appreciated that opioid analgesics elicit bimodal effects on nociceptor excitability, producing excitation as well as inhibition, *in vitro* ^10^. In behavioral studies we have previously shown that hyperalgesia induced by a systemic low, sub-analgesic dose of morphine, in rats, is nociceptor TLR4-dependent ^2^. Moreover, it has recently been reported that the mu-opioid analgesic, remifentanil, upregulates TLR4 expression in DRGs in the rat and decreases mechanical nociceptive threshold, an effect that is reversed by TAK-242, a TLR4 antagonist ^38^. In fact, activation of TLR4 produces robust mechanical and thermal hyperalgesia ^4, 14, 48, 55^, and many opioids at ultra-low doses are TLR4 agonists ^2, 23^. Since sub-analgesic doses of opioids produce TLR4-dependent hyperalgesia ^2, 23, 39^, hyperalgesia induced by the repeated administration of opioids (OIH) could be due to desensitization of MOR (i.e., opioid tolerance), uncovering TLR4-dependent hyperalgesia. In addition to direct activation of TLR4 by opioids, another mechanism by which TLR4 could contribute to OIH involves crosstalk between MOR and TLR4, such that opioids could activate TLR4 by acting at MOR ^2, 26, 57^. Both of these mechanisms implicate MOR in OIH, and a role for MOR is supported by our observation that OIH induced by a low systemic dose of morphine is partially dependent on MOR-mediated signaling (Fig. 2A in ^16^). However, the complete prevention of low dose morphine induced OIH by knock-down of TLR4 (Fig. 1B in ^2^) supports the suggestion that the switch from an inhibitory to an excitatory pathway for MOR signaling is unlikely to be a significant contributor to OIH.

In the present experiments, we have used well-established *in vitro* patch-clamp electrophysiology methods to study the effect of low dose morphine on excitability of small-diameter rat and human DRG neurons (putative C-type nociceptors) ^7, 15, 19, 21, 46, 54, 58^. Importantly, nociceptor excitability correlates with subjective pain levels in humans ^1, 27, 32, 41–43, 49^. In the current experiments we observed a similar sensitization (reduction of rheobase) of small-diameter neurons cultured from both rat and human DRGs, induced by low dose morphine. In addition, we observed that this morphine-induced nociceptor sensitization, *in vitro*, is dependent on the action of morphine on primary afferent nociceptor TLR4. This TLR4 dependence of nociceptor sensitization by morphine is based on the finding that pretreatment with a selective TLR4 antagonist (LPS-RS Ultrapure) prevents the sensitization of nociceptors by low dose morphine. These *in vitro* findings are consistent with our *in vivo* observation of the complete TLR4-dependence of OIH induced by a low dose of systemic morphine ^2^.

An excitatory effect of an opioid on human nociceptors may result in an attenuation of inhibitory effects of opioids, thereby reducing their analgesic efficacy, and necessitating greater doses to achieve adequate pain relief. This, in turn, increases the risk of other opioid side effects (e.g., nausea, constipation, addiction, impaired cognition, etc.) and facilitates the onset of tolerance. Although in the present experiments we evaluated a sub-analgesic dose of opioid, such an excitatory mechanism is likely to be also active at higher (analgesic) doses, but is masked by a stronger inhibitory effect. Developing tolerance to the MOR-dependent analgesia could, however, unmask a TLR4-dependent hyperalgesic action of opioids to result in OIH, considered a serious adverse effect of opioid analgesics. On the other hand, potentiation of a TLR4-dependent excitatory pathway (e.g., due to upregulation of TLR4 or enhanced crosstalk between MOR and TLR4), could also contribute to opioid tolerance as reduction in analgesic response to the same dose of opioid. Thus, our findings provide support for the hypothesis that using TLR4 antagonists in combination with opioid analgesics might increase the analgesic potency of opioids, as well as reduce risk and/or severity of TLR4-mediated opioid side-effects.

In conclusion, in the present study, we have validated the observation that opioid-induced nociceptor sensitization occurs by its direct action on small-diameter human and rat DRG neurons, enhancing the clinical relevance of prior *in vivo* and *in vitro* studies that have, to date, been conducted only in rat and mouse nociceptors. In addition, the present study provides strong support for the suggestion that opioid-induced nociceptor sensitization (and, by extension, OIH) is dependent on nociceptor TLR4, rather than indirectly by its action on other cells, such as cells of the immune system and glia.

## Acknowledgements

The authors thank Dionéia Araldi for her exclusive role in establishing contact with the provider of human tissue (AnaBios) and Paul G. Green for sharing his critical feedback and assistance in preparation of the manuscript. This study was funded by National Institutes of Health (NIH) grant R01 AR075334.

